# Neuron-specific transcriptomic signatures indicate neuroinflammation and altered neuronal activity in ASD temporal cortex

**DOI:** 10.1101/2022.03.29.486259

**Authors:** Pan Zhang, Alicja Omanska, Bradley P. Ander, Michael J. Gandal, Boryana Stamova, Cynthia M. Schumann

## Abstract

Autism spectrum disorder (ASD) is a highly heterogeneous disorder, yet transcriptomic profiling of bulk brain tissue has identified substantial convergence among dysregulated genes and pathways in ASD. However, this approach lacks cell-specific resolution. We performed comprehensive transcriptomic analyses on bulk tissue and laser-capture microdissected (LCM) neurons of 59 postmortem human brains (27 ASD and 32 matched controls) in the superior temporal gyrus (STG) ranging from 2-73 years of age. In bulk tissue, synaptic signaling, heat shock protein-related pathways and RNA splicing were significantly altered in ASD. There was age-dependent dysregulation of genes involved in GABA (*GAD1* and *GAD2*) and glutamate (*SLC38A1*) signaling pathways. In LCM neurons, AP-1 mediated neuroinflammation and insulin/IGF-1 signaling pathways were upregulated in ASD, while mitochondrial function, ribosome and spliceosome components were downregulated. GABA synthesizing enzymes *GAD1* and *GAD2* were both downregulated in ASD neurons. Alterations in small nucleolar RNAs (snoRNAs) associated with splicing events suggested interplay between snoRNA dysregulation and splicing disruption in neurons of individuals with ASD. Our findings supported the fundamental hypothesis of altered neuronal communication in ASD, demonstrated that inflammation was elevated at least in part in ASD neurons, and may reveal windows of opportunity for biotherapeutics to target the trajectory of gene expression and clinical manifestation of ASD throughout the human lifespan.

## Introduction

Autism spectrum disorder (ASD) defines a heterogeneous set of complex neurodevelopmental disorders affecting 1 in 54 children in the United States according to current estimation (1, 2) and confers lifelong challenges. ASD is characterized by difficulties with social communication as well as a repetitive, restricted repertoire of behaviors and interests (3). Population, family and twin studies all indicate a strong genetic component contributing to risk for ASDs (4, 5), with heritability estimates of ∼ 70% (6). However, the genetic causes and pathophysiology of ASD are varied and often complex.

Despite this heterogeneity, transcriptomic analyses of postmortem human brain have elucidated substantial convergent molecular-level pathology associated with idiopathic and syndromic forms of ASD (7-13). Multiple studies have profiled the transcriptomes of postmortem brain regions from individuals diagnosed with ASD (7, 8, 11, 14), including the temporal cortex implicated due to its critical importance in speech and language function (7, 8, 11). The most consistent findings include disruption of neuronal/synaptic activity and activation of innate immunity/glial markers (7, 8, 11). Dysregulation of alternative splicing and non-coding RNAs has also been shown to be dysregulated in ASD brains (8).

Most previous transcriptomic studies, however, profiled homogenate brain tissue and have therefore been unable to pinpoint the underlying specific cell-types in which gene expression is altered. Recently, Velmeshev *et al.* have published the first single-nucleus RNA-sequencing (sn-RNAseq) dataset in postmortem ASD cortex (13), identifying substantial changes in upper-layer excitatory neurons and microglia, consistent with observations from bulk tissue. As such sn-RNAseq datasets currently profile only the 3’ end of highly expressed genes within each cell, these data characterize neither lowly expressed coding and non-coding genes, nor splicing alterations that may contribute to altered neuronal function in ASD.

Here, we performed the first systematic study using transcriptomic profiling to directly compare both bulk cortical tissue and laser capture microdissected (LCM) neurons from anatomically well-defined superior temporal gyrus (STG) samples from 59 subjects (27 with ASD and 32 age-matched controls) ranging from 2-73 years of age (Figure 1, Supplementary Table 12). The STG modulates language processing and social perception, thereby playing a critical role in integrating a breadth of information to provide meaning to the surrounding world (15). Structural and functional imaging studies have long implicated STG in ASD (15, 16), however molecular-level changes in neurons remain unknown. This study aimed to identify neuron-specific transcriptomic changes in ASD brain by identifying differentially expressed genes, differential splicing events, age-related gene expression changes across the lifespan, as well as co-expression networks to reveal gene modules altered in ASD (Figure 1).

**Figure 1.**
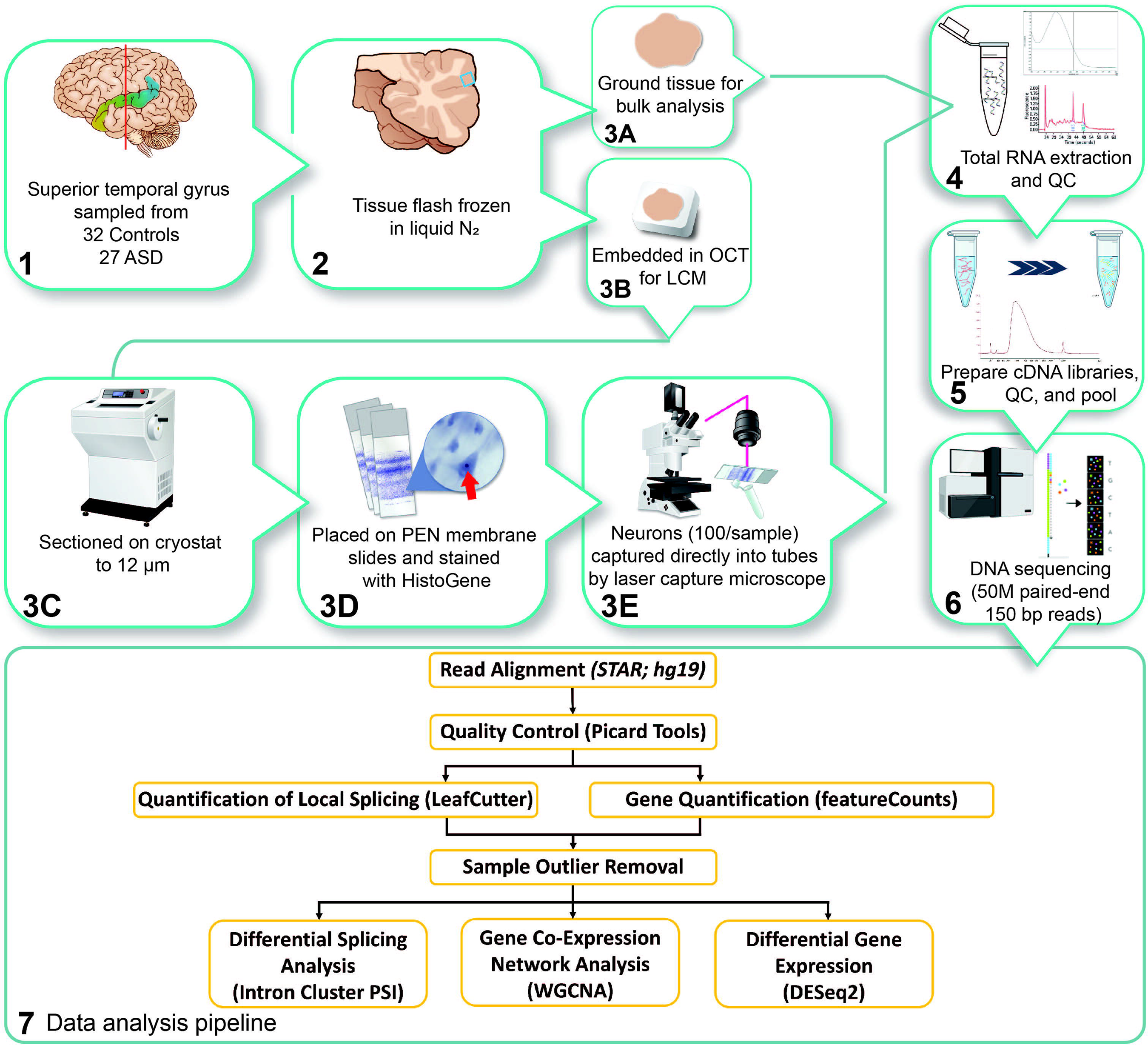
Overview of experiment design and data analysis pipeline.

## Results

### Global gene expression changes in ASD superior temporal gyrus (STG)

RNA sequencing was performed on bulk tissue STG of 59 human brains, 27 from individuals with ASD and 32 from age-matched neurotypical controls, ranging from 2-73 years of age. Following quality control, we performed a comprehensive characterization of differential gene expression and local splicing alterations in ASD. After adjusting for known covariates and correcting for multiple comparisons, we found 194 differentially expressed genes between individuals with ASD and controls (FDR < 0.05). Of these, 143 were upregulated and 51 were downregulated (Figure 2A, Supplementary Table 1), with a median absolute fold change of 1.45 (range 1.11 -4.04, Figure 2A). We observed significant concordance between our differential gene expression (DGE) results and previous data of the same region from different samples (12) (Supplementary Figure 1, Spearman ρ = 0.37 for t statistics, p-value < 10^−16^).

**Figure 2.**
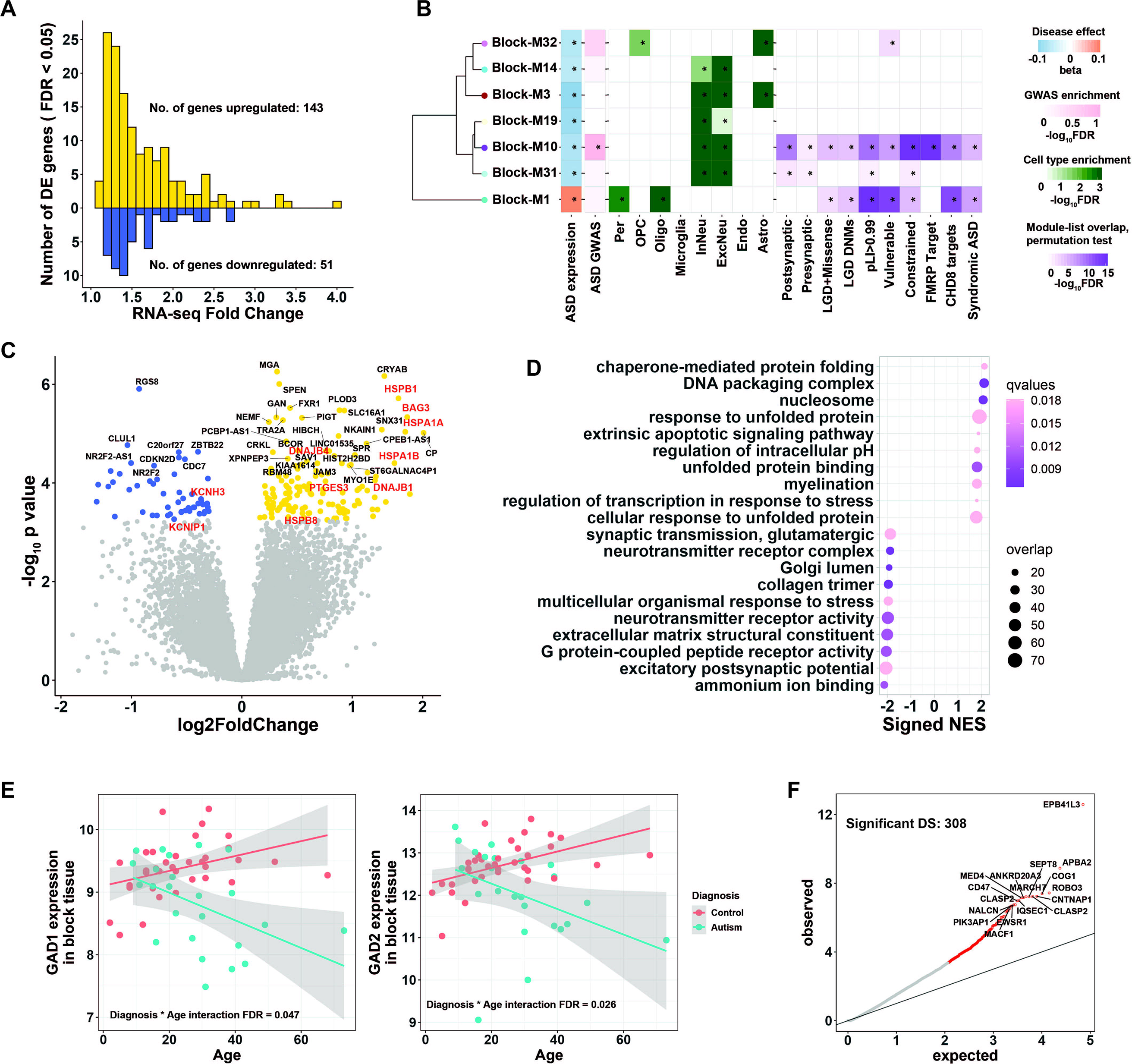
Transcriptomic difference between ASD cases and controls in bulk tissue STG. **A**, Distribution of fold-change of differential expression for 194 differentially expressed genes. Case:control fold-changes for upregulated genes are plotted in gold (N = 143, positive values) and control:case fold-changes for downregulated genes in blue (N = 51, negative values). **B**, Co-expressed gene modules that were significantly disrupted in ASD. Modules were hierarchically clustered by module eigengene. Module-diagnosis associations were shown on the right of each module (*FDR < 0.05). Additional enrichment analyses were also shown for each module, including: Enrichment for ASD GWAS common variants (30)(*FDR < 0.1); Enrichment for major CNS cell types(76)(*FDR < 0.05); Enrichment against literature-curated gene lists (*FDR < 0.05) including pre- and postsynaptic marker genes (77), genes with likely-gene-disruption (LGD) or LGD plus missense de novo mutations(DNMs) found in patients with neurodevelopmental disorders(78), genes with probability of loss-of-function intolerance (pLI) > 0.99 as reported by the Exome Aggregation Consortium (28), mutationally constrained genes(27), vulnerable ASD genes(79), CHD8 targets(29), FMRP targets (80), syndromic and highly ranked (1 and 2) genes from SFARI Gene database. Abbreviations: Per, pericytes; OPC, oligodendrocyte progenitor cells; InNeu, inhibitory neuron; ExcNeu, excitatory neuron; Oligo, oligodendrocytes; Endo, endothelial cells; Astro, astrocytes. **C**, Volcano plot showing significantly up- (gold) and down-regulated (blue) genes (FDR < 0.05). Genes discussed in the main text are colored red. **D**, Functional enrichment of differentially expressed genes in ASD cases compared to controls. Top ten significantly enriched up- and down-regulated categories were shown. Categories were ranked by normalized enrichment score (NES), and NES for down-regulated categories were set to negative for displaying purpose solely. The color of each dot reflects FDR-corrected q-value, and the size of each dot reflects the number of overlapped genes between our gene list and the corresponding GO category. **E**, Age trajectory of GAD1 (left) and GAD2 (right) gene expression, stratified by ASD diagnosis. **F**, Quantile-quantile plot of observed p-values vs expected p-values for differentially spliced intron clusters. Significant DS events (FDR < 0.05) were colored red, and overlapping gene names were labeled for top clusters.

Functional and pathway enrichment analyses indicated an over-representation of heat shock proteins (HSPs) and HSP-related chaperones, which were upregulated in ASD subjects. This included HSP70 family members *HSPA1A* and *HSPA1B*; HSP40 family members *DNAJB1* and *DNAJB4*; small HSP20 family members *HSPB1* and *HSPB8*; and HSP-binding chaperons *BAG3* and *PTGES3* (Figure 2C,D). HSPs are involved in stress-response, immune activation and inflammation(17, 18), all of which were upregulated in ASD postmortem brain (7). Downregulated genes were mainly enriched in pathways related to synaptic function (Figure 2D), consistent with previous findings (7). Notably, two important voltage-gated potassium channel-related genes *KCNH3* and *KCNIP1* were among the most downregulated (Figure 2C), which may relate to disrupted neuronal excitability hypothesized in ASD (19, 20).

As age-dependent expression alterations have been reported in ASD brain (14), we employed an analytical model accounting for age and the interaction between age and diagnosis. Fourteen genes showed age-dependent DGE between ASD and control (Supplementary Table 2). Interestingly, genes involved in gamma aminobutyric acid (GABA) synthesis (*GAD1* and *GAD2*) (21) were downregulated in ASD only during late adulthood (Figure 2E). This may indicate an age-dependent dysregulation of GABA signaling in ASD neurons, or a decrease in the proportion of GABAergic neurons in ASD brains (22).

Differential splicing (DS) events in the bulk tissue transcriptome were evaluated using LeafCutter (26). Among 35,505 intron clusters identified by LeafCutter, 308 clusters (297 unique genes) showed significant DS between ASD patients and controls (FDR < 0.05). The 297 genes did not show significant functional enrichment (Figure 2F, Supplementary Table 5).

To place subtle changes across the ASD STG transcriptome into a systems-level context, we performed weighted gene correlation network analysis (WGCNA) to build gene co-expression networks (23), identifying 31 modules of co-expressed genes (Methods; Supplementary Table 3). Seven modules showed a significant association with ASD diagnosis, two of which were strongly enriched for ASD-associated genetic risk factors (Modules Block-M1 and Block-M10, Figure 2B).

Module Block-M1 was upregulated in ASD STG and its gene members were enriched in RNA splicing and mRNA metabolic pathways (Supplementary Table 4). Notably, significantly upregulated HSPs were also members of the Block-M1 module (Supplementary Table 3). HSPs contribute to RNA splicing during stress (24). Downregulated modules in ASD were mostly enriched for synaptic functions (Block-M3, Block-M10, Block-M14, Block-M19, Block-M31; Supplementary Table 4). Cell-type enrichment analysis also indicated these downregulated modules were enriched in marker genes for both excitatory and inhibitory neurons (Figure 2B), suggesting a broad disruption of neuronal and synaptic processes in ASD STG.

Genes in the upregulated module Block-M1 and one downregulated module (Block-M10) were enriched in high-confidence ASD risk loci (25, 26), mutationally constrained (27) and highly intolerant to mutations (pLI > 0.99) genes (28), as well as regulatory target genes of CHD8, which has clear links to at least a subset of ASD cases (29) (Figure 2B). Many hub genes for module Block-M10 encoded synaptic proteins (Supplementary Figure 2, Supplementary Table 3). This module was also enriched for ASD common risk alleles from ASD GWAS data (30). Together this suggested a causal role of synaptic dysfunction in ASD etiology.

### Neuron-specific gene expression and splicing alterations in ASD STG

To provide cell-type specificity for the observed transcriptomic changes, we next performed laser capture microdissection to capture neurons using STG sections taken from the same subjects profiled using bulk RNA-seq. We then interrogated ASD-associated gene expression and splicing alterations using the same bioinformatic pipelines as above. Across 13,458 neuron-expressed genes, 83 were significantly differentially expressed between ASD subjects and controls at FDR < 0.05, of which 52 were upregulated and 31 downregulated (Figure 3A, Supplementary Table 6). Median absolute fold change in expression between subjects with ASD and controls was 2.48 (range 1.29 - 9.72; Figure 3A). Surprisingly, concordance of neuronal DGE with bulk tissue DGE was low (Spearman ρ = 0.18 for t statistics, Supplementary Figure 3), suggesting our analysis captured ASD signatures unique to neurons.

**Figure 3.**
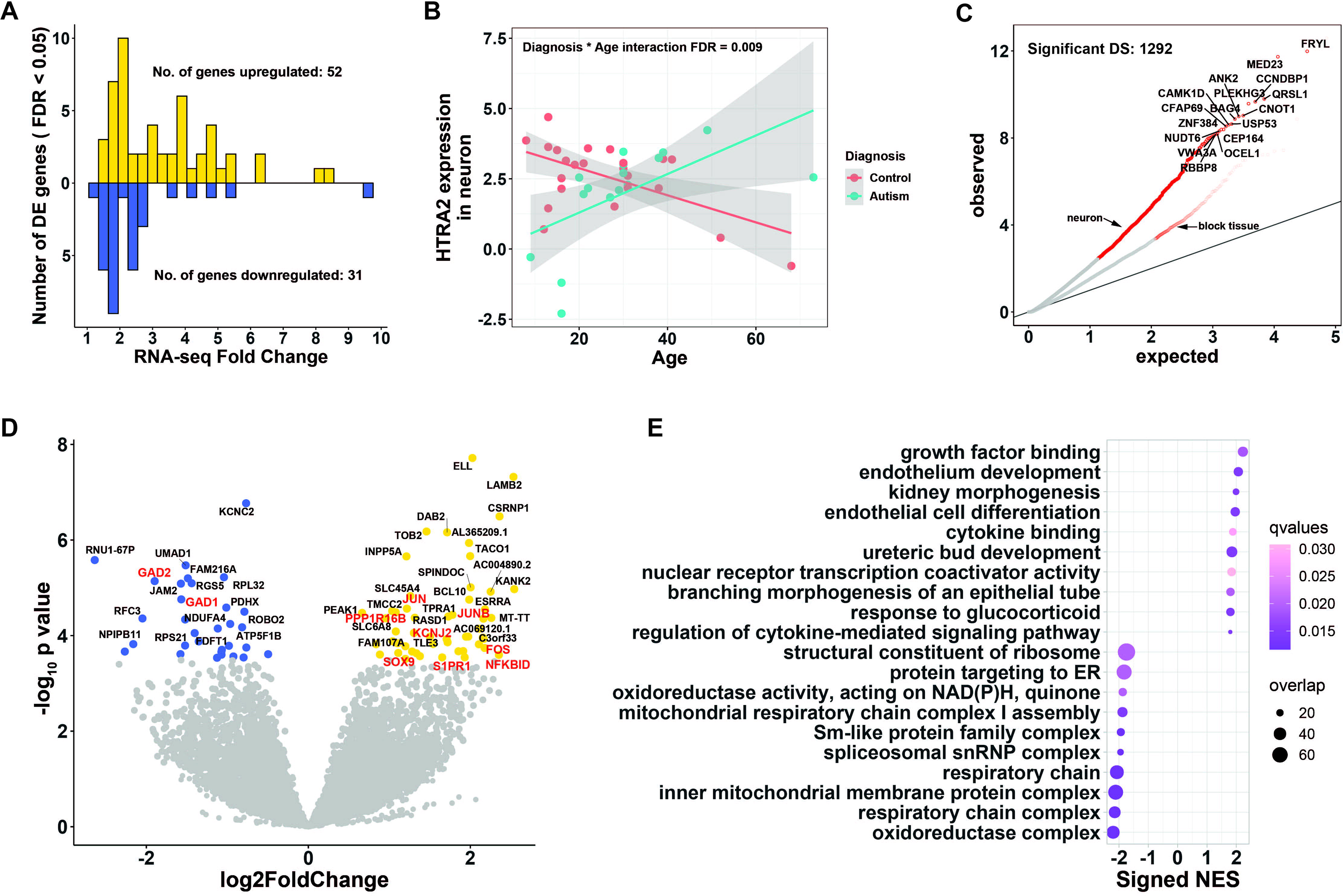
Differential gene expression and differential splicing between ASD cases and controls in LCM neurons from STG. **A**, Distribution of fold-change of differential expression for 83 differentially expressed genes. Case:control fold-changes for upregulated genes are plotted in gold (N = 52, positive values) and control:case fold-changes for downregulated genes in blue (N = 31, negative values). **B**, Age trajectory of HTRA2 gene expression, stratified by ASD diagnosis. **C**, Quantile-quantile plot of observed p-values vs expected p-values for differentially spliced intron clusters. Significant DS events (FDR < 0.05) were colored red, and overlapping gene names were labeled for top clusters. DS results from bulk tissue were also plotted for comparison. **D**, Volcano plot showing up- (gold) and down-regulated (blue) genes. Genes discussed in the main text are colored red. **E**, Functional enrichment of differentially expressed genes in ASD cases compared to controls. Top ten significantly enriched up- and down-regulated categories are shown. Categories were ranked by normalized enrichment score (NES), and NES for down-regulated categories were set to negative for display purposes solely. The color of each dot reflects FDR-corrected q-value, and the size of each dot reflects the number of overlapped genes between our gene list and the corresponding GO category.

Upregulated genes in ASD neurons were highly enriched in pathways related to growth and differentiation (Figure 3E). Specifically, the AP-1 transcription factor complex components (31) *FOS*, *JUN* and *JUNB* were all up-regulated in ASD neurons, among other growth/differentiation regulators such as *SOX9*, *S1PR1* and *PPP1R16B* (Figure 3D). The AP-1 transcription factor complex is known to regulate a number of downstream biological processes (31). Indeed, we found upregulated B cell signaling adaptor gene *BCL10* and NFκB inhibitor delta gene *NFKBID* in ASD neurons (Figure 3D). NFκB and AP-1 may function together in regulating inflammatory processes (32-34), and up-regulation of both *NFKBID* and AP-1 point to dysregulated inflammation in ASD neurons. In addition, the inward-rectifier potassium ion channel gene *KCNJ2* was also up-regulated in ASD neurons (Figure 3D). Interestingly, both *KCNJ2* and AP-1 subunit *FOS* were involved in regulating excitability and plasticity at the cholinergic synapse (35-37).

Downregulated genes in ASD neurons are primarily enriched in mitochondrial function and oxidoreductase activity (Figure 3E). Specifically, comparing to bulk tissue STG, more subunits of the NADH:ubiquinone oxidoreductase (complex I) were downregulated in neurons, and their effect sizes were larger (Supplementary Figure 4). Indeed, oxidative phosphorylation (OXPHOS) defects have been reported in ASD lymphocytes, muscle, and temporal lobe (38-40). Our results provide evidence that compared to bulk tissue, mitochondrial dysfunction is much more profound in STG neurons.

While LCM captured both excitatory and inhibitory neurons, we note that *GAD1* and *GAD2* genes are among the most downregulated in ASD neurons (Figure 3D). The coordinated down-regulation of both GABA synthesizing enzymes suggest that the level of GABA neurotransmitter may be decreased in ASD neurons, providing support to the excitation to inhibition (E/I) imbalance hypothesis of ASD (19, 20, 41).

By testing the interaction between diagnosis and age, 3 genes (*HTRA2,* aka *OMI*; *ZNF765*; and *PCDHB18P*) showed age-dependent differential expression in ASD neurons (Supplementary Table 7). For example, age trajectories of serine peptidase *HTRA2* were opposite in ASD brains compared to controls (Figure 3B). In healthy brains, the expression of *HTRA2* was much higher before age 30 and decreases with age, while its expression levels begin lower and increase with age in ASD STG neurons. Attenuated *HTRA2* activity may lead to neuronal cell death, altered chaperon activity and autophagy and has been linked to Parkinson’s disease (42). In addition, increased active form of the *OMI/HTRA2* serine protease has been positively correlated with cholinergic alterations in AD brain (43). Thus, it is plausible that the altered expression of *HTRA2* with age we observed in ASD brain may be associated with neuronal alterations during development.

We also quantified local splicing events in the neuronal transcriptome. After adjusting for multiple testing, LeafCutter identified 1292 significant differential spliced intron clusters (1177 unique genes) out of 17,250 total intron clusters at FDR < 0.05 (Figure 3C, Supplementary Table 8). No functional enrichment was observed for the 1177 genes. We observed more disruptions in local splicing events in ASD neurons than in bulk tissue (308 DS out of 35505 events in bulk tissue, 1292 DS out of 17250 events in neurons; p < 2 x 10^-16^ test of proportions).

### Neuron-specific networks pinpoint subtle changes in the neuronal transcriptome in ASD

Co-expression network analysis on neuronal data identified 18 modules, each containing between 101 and 998 co-expressed genes (Supplementary Table 9). Four modules were significantly upregulated in ASD neurons, while one module was downregulated (Figure 4A).

**Figure 4.**
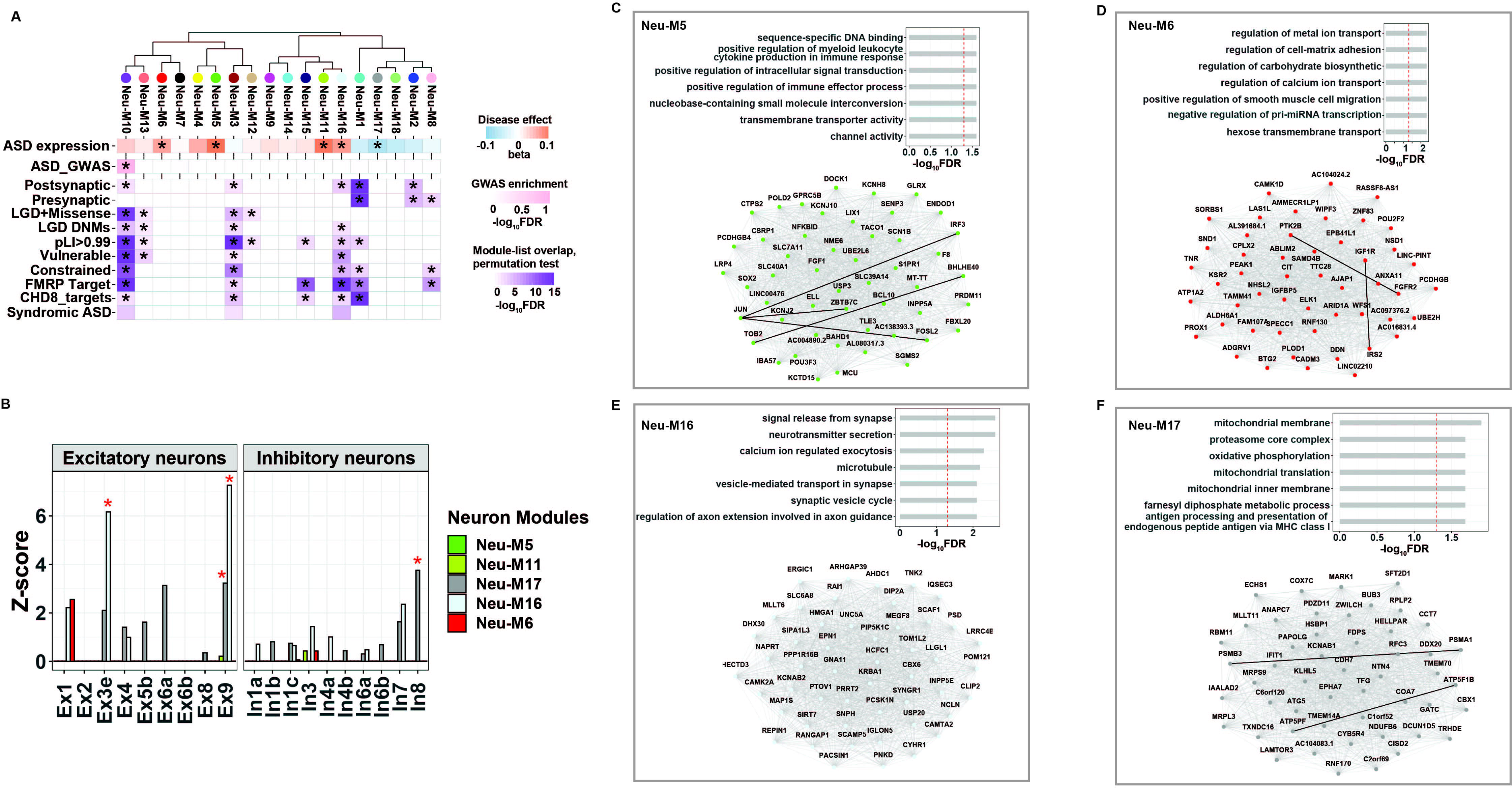
Gene co-expression network analysis of the ASD neuronal transcriptome. **A**, Hierarchical clustering of neuronal gene co-expression modules by module eigengenes. Module-diagnosis associations were shown below each module. Enrichment for ASD GWAS common variants is shown for each module. Enrichment against literature-curated gene lists is shown on the bottom. **B**, Module enrichment for neuron subtypes. Expression profiles of neuron subtypes were obtained from ref. (76). Red asterisks indicate significant enrichment. **C-F**, Functional enrichment (top panel) and top 50 hub genes (bottom panel) for module neu-M5 (C), neu-M6 (D), neu-M16 (E) and neu-M17 (F). Edges represent co-expression (Pearson correlation > 0.5). Co-expressed partners with evidence of protein-protein interaction were connected by solid black lines. PPI data was compiled from well-characterized PPI databases, including Bioplex, HPRD, Inweb, HINT, Biogrid, GeneMANIA, STRING and CORUM. Only physical interactions and co-complex associations were kept.

The upregulated neu-M5 co-expression module was highly represented by the DGE analysis signal. Upregulated genes *JUN, JUNB, NFKBID* were all hub genes of neu-M5 module. Neu-M5 module also captured additional AP-1 subunits and interactors, such as *FOSL2* (44) and *IRF3* (34, 45). Neu-M5 module was enriched in immune response pathways, providing further evidence that AP-1 mediated neuroinflammation was elevated in ASD neurons (Figure 4C, Supplementary Table 10). Hubs of Neu-M5 also contained multiple ion channel-related genes, such as sodium ion channel gene *SCN1B*, potassium channel genes *KCNJ2* and *KCNJ10*, and solute carrier gene *SLC40A1* (Figure 4C, Supplementary Table 9). Coordinated upregulation of various ion channels suggested that membrane transport was activated in ASD neurons, consistent with heightened excitability. Neu-M5 was significantly overlapped with module M16 from *Voineagu et al. (7)*. M16 was also enriched in immune/inflammatory response and was up-regulated in ASD (7). Our data refined our understanding of the neuroinflammatory changes in ASD to include a neuronal component. Additionally, downregulated neu-M17 module was enriched in mitochondrial function and contained most differentially expressed mitochondrial genes, such as ATP synthase subunits *ATP5F1B* and *ATP5PF* (Figure 4F, Supplementary Table 9).

Neuronal co-expression networks further captured signals that were not detected by DGE analysis. Neu-M6 module was upregulated in ASD, and among its hub genes were several insulin signaling pathway components, including insulin-like growth factor (IGF) receptor *IGF1R*, IGF binding protein *IGFBP5*, insulin receptor substrate *IRS2* as well as CBL-associated *SORBS1* (Figure 4D, Supplementary Table 9). Insulin signaling is associated with multiple neurodevelopmental disorders, including monogenetic ASD syndromes such as Rett and Phelan-McDermid syndromes (46-49). Our results provided direct molecular-level evidence that insulin signaling was altered in ASD neurons.

Among all five significantly disrupted modules, none were enriched for ASD common variants and only one upregulated module (Neu-M16) showed enrichment in highly confident ASD risk genes, as well as in several other curated gene sets (Figure 4A). Neu-M16 was enriched in synaptic functions (Figure 4E, Supplementary Table 10). Further, cell-type analysis showed that Neu-M16 was also highly enriched in excitatory neurons (Figure 4B), with *CAMK2A* and *CAMK2B* among its hub genes (Supplementary Table 9). The upregulation of Neu-M16 suggested elevated excitatory signal in ASD neurons.

We further tested if significantly disrupted neuronal modules were enriched in any neuron subtypes. Upregulated neuronal modules in ASD were only enriched in excitatory neuron subtypes while enrichment of inhibitory neurons was only observed for downregulated modules (Figure 4B). This provides additional evidence for altered neuronal activity in ASD neurons, consistent with the findings in our DGE analysis.

### Small non-coding RNAs are selectively down-regulated in ASD neurons and correlate with altered local splicing

When investigating the genes downregulated in ASD neurons more closely, we noticed a striking pattern that, 51 out of the 59 neuron-expressed small nucleolar RNA (snoRNA) (50) genes were down-regulated in ASD neurons, and 13 were significantly down-regulated at p-value < 0.05 (Figure 5A, Supplementary Table 6). Dysregulation of snoRNAs was not observed in bulk tissue (Supplementary Table 1), and snoRNAs were undetectable in a recent ASD single-cell study (13). snoRNAs are involved in the modification and maturation of ribosomal RNAs (rRNAs) and small nuclear RNAs (snRNAs) (51). Interestingly, both ribosome and spliceosome components were among the most downregulated in ASD neurons (Figure 3E). Moreover, snRNAs were also downregulated in ASD neurons, with 23 out of 24 snRNA genes down-regulated in ASD and 13 significantly down-regulated at p-value < 0.05 (Supplementary Figure 5, Supplementary Table 6).

**Figure 5.**
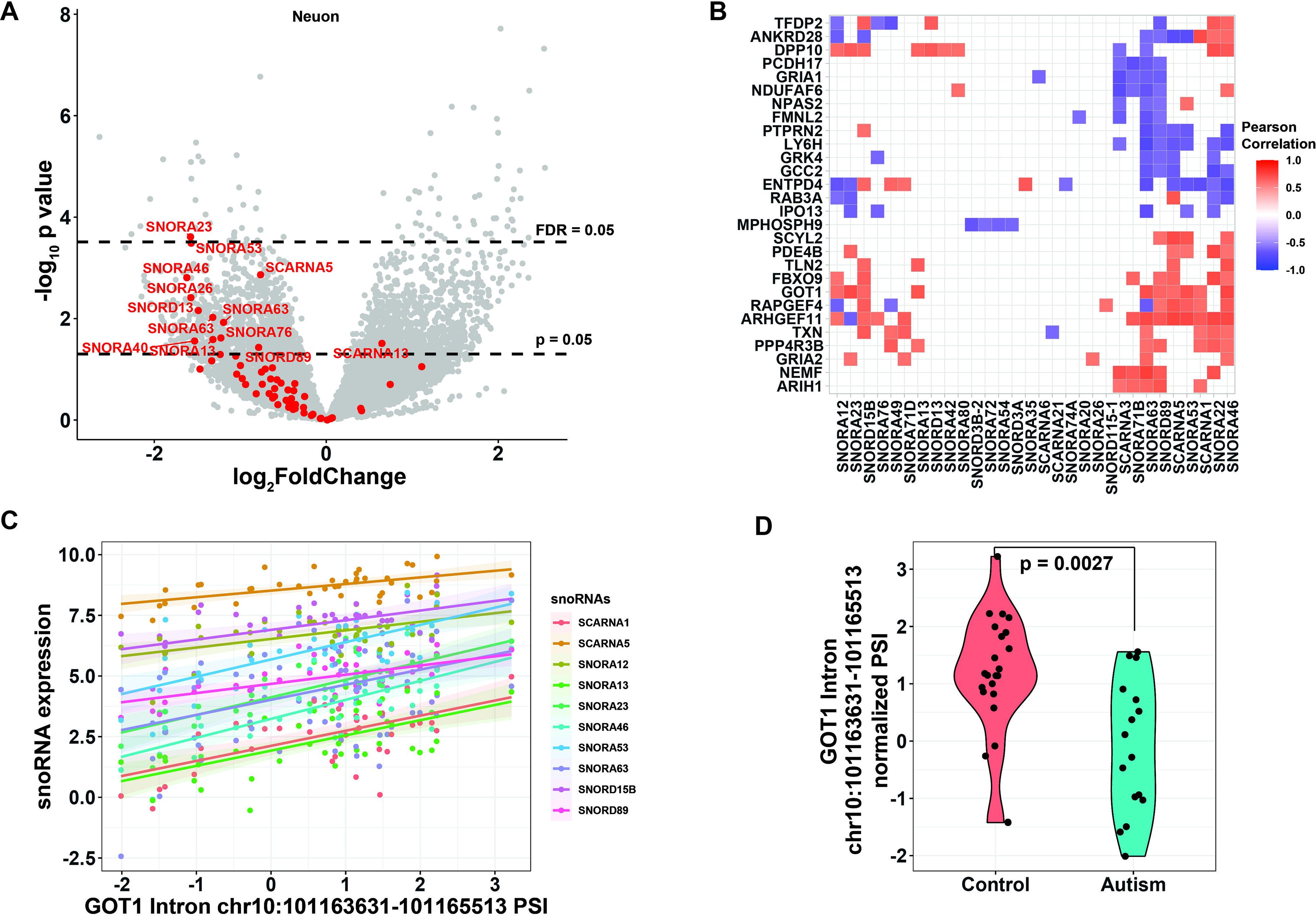
Coordinated dysregulation of snoRNAs in ASD neurons. **A**, Volcano plot showing differentially expressed genes in ASD neurons compared to control. snoRNA genes were colored red. **B**, Significant correlation between snoRNA expression (x-axis) and intron PSI (y-axis). Introns are labeled with the name of overlapping gene locus. Gene loci that are correlated with more than 3 snoRNAs were shown. **C**, Scatter plot showing the correlation between GOT1 intron 5 PSI and expression levels of multiple snoRNAs across all neuron samples. Also shown were fitted regression lines with 95% confidence intervals. Intron coordinates were based on GRCh37. **D**, PSI of GOT1 intron 5 is downregulated in ASD neurons.

As snoRNAs and snRNAs are known to be critical regulators of alternative splicing (52-55), and splicing alterations are strongly implicated in ASD and observed in LCM-captured neurons, we next examined alterations in splicing events that may be associated with snoRNA dysregulation. We calculated the correlation between snoRNA expression level and each local splicing event in neuronal data, followed by FDR correction for multiple comparisons. We identified 835 gene loci in neurons with at least one intron whose percent spliced in (PSI) was significantly correlated with snoRNA gene expression (Supplementary Table 11). Of these 835 intron clusters, 196 were significantly dysregulated in ASD neurons (Supplementary Table 11). Several intron clusters correlated with multiple snoRNAs and corresponded to genes involved in synaptic functions (Figure 5B). For example, *GOT1* encodes the glutamic-oxaloacetic transaminase known to function as an important regulator of glutamate level (56). PSI of intron 5 of *GOT1* was highly correlated with the expression level of multiple snoRNA genes (Figure 5C). *GOT1* intron 5 was also differentially spliced between ASD and control (Figure 5D). Differential splicing of *GOT1* gene may change the level of glutamate, and thus leads to an imbalance of E/I in neuronal communication in ASD neurons.

## Discussion

Altered neuronal processes and synaptic function are consistent findings in transcriptomic analyses of ASD brain (7, 8, 11). However, transcriptomic studies of autistic brains are mostly limited to bulk tissue (7, 8, 11). Cell-type-specific molecular alterations in neurons are still largely unexplored. A recent single-cell RNA-seq study sheds light on the cell-type-specific transcriptomic changes in ASD (13), however, due to technology limitations, single-cell studies currently cannot reliably quantify low-expressed genes and local splicing events. Thus, in this study, we performed comprehensive analyses of gene expression and alternative splicing by short-read paired-end RNA-seq on both bulk tissue and LCM-isolated neurons from the STG brain region in a cohort of 59 human brains ranging from 2-73 years of age.

### Bulk tissue transcriptome findings reveal downregulated neuronal and synaptic function processes, upregulation of heat shock proteins, and unfolded protein response in ASD temporal cortex

Our bulk tissue analyses revealed a potential causal role of downregulated neuronal processes and synaptic functions in ASD etiology, consistent with findings from previous bulk-tissue transcriptomic studies on the same brain region (7). Previous studies also reported dysregulated alternative splicing events in ASD brain (8, 57). Differential splicing analysis in bulk STG found several synaptic genes, including *CANCA2D1*, *CAMK4*, *CLASP2*, *CNTNAP1*, *EPHB1*, *KALRN*, *NRXN3*, *SOS2* and *SYNGAP1*, differentially spliced in ASD. *SYNGAP1* isoforms have been shown to differentially regulate synaptic plasticity and dendritic development (58). This further signifies the importance of studying alternatively spliced isoforms in ASD brain.

We also observed a coordinated upregulation of multiple HSPs and HSP-related chaperones in ASD STG. HSPs can serve as activators and regulators of the immune system (18), and upregulated HSPs may induce immune responses in ASD brain. They also play a role in facilitating alternative RNA splicing (24). Previous studies found that both immune response and RNA splicing are upregulated in ASD brain (7, 8, 11), and our results signify that upregulated HSP-related pathways are a potential contributor to these observations. HSPs and HSP-related chaperones are normally induced in response to stress. The upregulation of HSPs in ASD neurons may relate to elevated endoplasmic reticulum (ER) stress since both unfolded protein response (UPR) and apoptosis are also upregulated in our ASD bulk data (Figure 2D). ASD-linked rare or *de novo* mutations in synaptic genes can lead to misfolded proteins and cause ER stress (59), itself coupled to heightened inflammation and neurotoxic cell death (60). ER stress-related genes are also dysregulated in the middle frontal cortex of subjects with ASD (61). Our data suggest that ER stress serves as a major response to ASD genetic mutations, and ER stress activates UPR, including the production of HSPs and chaperones. UPR further induces multiple downstream processes such as inflammation and immune response (62). Limiting the effect of ER stress and UPR may be a promising therapeutic avenue for ASD.

### ASD neuronal transcriptome reveals upregulated neuroinflammation and altered neuronal activity

We observed a strong upregulation of AP-1 transcription factor components in ASD neurons. AP-1 subunits *FOS*, *JUN* and *JUNB* were upregulated at FDR < 0.05, and *FOSL2* was upregulated at nominal p < 0.05. AP-1 regulated gene expression in response to a variety of stimuli, including cytokines, growth factors, stress signals, infections and inflammation/neuroinflammation (31). In ASD neurons, it is likely that AP-1 activation induces broad inflammatory response, since several immune and inflammation-related genes were also strongly upregulated. These included *NFKBID* and *BCL10*, both of which were involved in the NF-κB pathway and were upregulated at FDR < 0.05. In addition, the interferon regulatory factor *IRF3* is upregulated at p < 0.05, and *IRF3* is co-expressed with AP-1 subunits. The simultaneous upregulation of AP-1 subunits, NFκB-related genes and interferon regulatory factors suggested that immune and inflammation responses were activated in ASD neurons. Upregulated immune response and neuroinflammation have been consistently observed in ASD patients by bulk tissue transcriptomic studies largely implicating glial cells (7, 8). However, our results demonstrated that immune/neuroinflammatory response was clearly activated in ASD neurons, and may be mediated by transcription factor AP-1. In addition, analysis of the potential upstream regulators of the observed changes in the ASD neuronal transcriptome with Ingenuity Pathway Analysis predicted their activation. This includes MTORC2 member *RICTOR*, growth factors *FGF2* and *BMP4*, and *OSM* cytokine, most of which were implicated in ASD (63).

We also observed strong downregulation in ASD STG neurons of *GAD1* and *GAD2* genes, involved in the biosynthesis of the inhibitory neurotransmitter GABA. In contrast, *CAMK2A* and *CAMK2B* genes, which are essential for aspects of plasticity at glutamatergic excitatory synapses, are upregulated at nominal significance. In addition, co-expressed gene modules that were upregulated in ASD neurons were mainly enriched in excitatory neurons, while the downregulated module was primarily enriched in inhibitory neurons. These data provided further support for the hypothesis that ASD reflects imbalance of E/I in neuronal communication, also reported in several brain regions in ASD (19, 20, 41). To our knowledge this is the first report providing molecular-level evidence for imbalance of E/I in neuronal communication specifically in STG neurons in ASD.

Multiple insulin signaling pathway components, such as *IGFBP5*, *IRS2* and *SORBS1*, are coordinately upregulated in ASD neurons. Insulin signaling pathway is implicated in several neurodevelopmental disorders, likely due to its role in protein homeostasis and synaptic plasticity (46-49). Although the insulin-like peptide IGF-1 is currently in clinical trial for ASD, the direction of change in ASD brain is still controversial (49). In our data, the expression level of IGF-1 is downregulated in ASD neurons at nominal significance; upregulation of the aforementioned insulin signaling pathway components may reflect a compensatory reaction to the lack of IGF-1 ligand.

Future studies will focus on the role of snoRNAs in ASD neurons, as well as other long and small modulatory non-coding RNAs. Given the emerging role of snoRNAs as alternative splicing regulators (54, 55), we hypothesize that a coordinated downregulation of multiple snoRNAs correlates with elevated dysregulation of local splicing events in ASD neurons. Our data provide evidence supporting this hypothesis, however no causal relationship can be determined. It will be critical to determine if snoRNA dysregulation plays a causal role in ASD etiology, and if so, pinpoint the underlying mechanism and possibly relate to transcript isoforms driving ASD brain and neuronal phenotype.

### Age-associated differential expression in STG points to altered neuronal activity in ASD

Age-dependent gene expression changes have been observed in the ASD brains (64, 65). We previously reported age-dependent miRNA alterations in the superior temporal sulcus (STS) and adjacent primary auditory cortex (PAC) (64). In this study, a number of genes show varying age trajectories in ASD bulk tissue and isolated neurons.

In bulk tissue, genes involved in GABAergic signaling (*GAD1* and *GAD2*) were upregulated with age in controls, while downregulated with age in ASD. Multiple lines of evidence have pointed to reduced neuronal inhibitory signal as a hallmark of ASD, including decreased number of GABAergic interneuron (especially Parvalbumin neurons) (66) and reduced density of GABA receptors (67-70). These cellular phenotypes are mainly observed in adults. Our findings indicated that the reduction of *GAD1* and *GAD2* mRNA levels in ASD brain became more profound with increasing age, consistent with the observations at cellular level. In addition, *SLC38A1*, involved in neurotransmission at glutaminergic and GABAergic synapses (71), was downregulated in ASD relative to control STG. *SLC38A1* is implicated in Rett Syndrome (72) and mitochondrial disorders, and its decrease may contribute to the observed alterations in synapse formation and neural connectivity.

In LCM neurons, the expression of *HTRA2* was higher below age 30 and decreases with age in control neurons, while lower at younger ages and increasing with age in ASD neurons. HTRA2 is important in maintaining mitochondrial homeostasis (73) and inducing apoptosis. It is implicated in pathogenesis of neurodegeneration, hypoxic-ischemic damage, and is proposed as a potential treatment target in neurological diseases (74). These findings further support the hypothesis of altered neuronal E/I activity, neuroinflammation, cell death, and mitochondrial dysfunction, implicated in ASD (75) and suggest treatment windows to target specific genes to alter their expression trajectories with age.

### Closing

Our study examined both bulk cortical tissue and isolated neurons from STG of autism and control brain. We found expression patterns in neurons that are not detectable when aggregate transcriptomes of multiple cells are profiled. In both cases, transcriptomic evidence suggested alteration in E/I balance could contribute to asynchronous activity in the brain which may be a contributing factor to autism in early development. Neuronal transcriptomes revealed further and more specific evidence of divergent activity and molecular signaling pathways. Future studies will need to examine the roles of other cell types in the brain and their contribution to ASD phenotype. As research into ASD continues to focus on more precise data, it is apparent that the complexity of factors that produce an ASD phenotype are slowly becoming clearer. As recent technologies are applied to the valuable collections of banked brain tissue, better definitions of specific subsets of ASD will become possible.

## Figure Legend

**Supplementary Figure 1** Binned density scatter plot comparing the t-statistics for case versus control differential expression between this study and another study (De Bree *et al.*, unpublished) comparing gene expression between ASD and controls in BA41, BA42, and BA22 bulk tissues; correlation between the statistics is 0.37 (P < 10^−16^).

**Supplementary Figure 2** Representative genes in module Block-M10. Known ASD risk genes were colored red. Synaptic genes that are intolerant to LOF mutation were colored pink. Edges represent co-expression.

**Supplementary Figure 3** Binned density scatter plot comparing the t-statistics for case versus control differential expression between neurons and bulk tissue; correlation between the statistics is 0.18 (P < 10^−16^).

**Supplementary Figure 4** Fold changes (ASD *vs.* CTL) of NADH:ubiquinone oxidoreductase (complex I) subunits in block tissue (red) and neurons (green).

**Supplementary Figure 5** Volcano plot showing differentially expressed genes in ASD neurons compared to control. snRNA genes were colored red.

**Supplementary Table 1** DGE summary statistics for block tissue

**Supplementary Table 2** DGE summary statistics for age-diagnosis-interaction in block tissue

**Supplementary Table 3** Gene co-expression network module membership for block tissue

**Supplementary Table 4** Gene co-expression network module functional enrichment for block tissue

**Supplementary Table 5** DS summary statistics for block tissue

**Supplementary Table 6** DGE summary statistics for LCM neuron

**Supplementary Table 7** DGE summary statistics for age-diagnosis-interaction in LCM neuron

**Supplementary Table 8** DS summary statistics for LCM neuron

**Supplementary Table 9** Gene co-expression network module membership for LCM neuron

**Supplementary Table 10** Gene co-expression network module functional enrichment for LCM neuron

**Supplementary Table 11** Significant correlations between snoRNA gene expression and local splicing events

**Supplementary Table 12** Donor information

## Methods

### Block tissue RNA extraction and library preparation

Human brain tissue was collected, sectioned coronally and flash frozen. STG from 32 controls and 27 ASD cases (2-73 years old) was identified anatomically according to “Atlas of the Human Brain” 4th edition (Maj, Majtanik, Paxinos 2015). Brain tissue (18-25 mg) was excised from the STG and put directly into 600 μl of Tri Reagent lysis buffer. Total RNA was extracted using the Direct-zol RNA MiniPrep (Zymo Research #R2051) following manufacturer’s protocol, with the inclusion of DNase I treatment and eluted in DNase/RNase-free water. Quality and quantity of RNA were determined via RNA 6000 Nano chip on 2100 Bioanalyzer (Agilent), NanoDrop 2000 spectrophotometer (ThermoFisher Scientific), and Qubit fluorometer (ThermoFisher Scientific).

From each of the 48 STG samples, 50 ng of RNA were used to create strand-specific total RNA libraries with the NuGEN Ovation Universal RNA-Seq System v2 and processed in parallel on the Sciclone NGS automated workstation (Perkin Elmer) according to manufacturer protocol. Following second-strand cDNA synthesis, samples were sheared by sonication on the Covaris E220. InDA-C (aka AnyDeplete) primers were used to target and cleave adapters from rRNA transcripts before amplification of libraries through 16 cycles of PCR. Final barcoded libraries were bead purified and examined for QC using 2100 BioAnalyzer DNA High Sensitivity chips. Library concentration was calculated based on fragment size and normalized to 15 nM for sequencing.

### Laser capture microdissection, RNA extraction and library preparation

Fresh-frozen STG tissue samples from 22 controls and 18 ASD cases (8-73 years old) were carefully dissected and embedded in OCT compound. The specimens were sectioned on a Microm HM550 cryostat (Thermo Scientific) at 12 μm and mounted on PEN membrane slides (ThermoFisher Scientific #LCM0522). Sections were hydrated with an ice-chilled ethanol series (100%, 75%, 50%) for 2 min each followed by HistoGene staining solution (ThermoFisher Scientific #KIT0415) for 30 seconds, 2 rinses in nuclease-free water, and alcohol dehydration (50%, 75%, 95%, 100% with molecular sieves). Slides were air dried and maintained on dry ice until laser capture microdissection.

Using a Leica LMD-6000 laser capture microdissection system, 100 neurons from each sample were laser captured directly into lysis buffer (PicoPure RNA Isolation Kit, ThermoFisher Scientific KIT0204). RNA was extracted using the PicoPure total RNA kit with inclusion of DNase I digest according to manufacturer protocol.

Strand-specific rRNA depleted RNA libraries were prepared from 10 μl of the final neuronal RNA eluate using the NuGEN Ovation SoLo Kit (NuGen #0500) for ultra-low input following manufacturer protocol with final amplification of 18 PCR cycles. Barcoded bead purified libraries were examined for QC using Qubit Fluorometer (ThermoFisher Scientific) and 2100 BioAnalyzer DNA High Sensitivity chips. Library concentration was calculated based on fragment size and normalized to 15 nM for sequencing.

### RNA sequencing

Library concentrations were confirmed with qPCR and pooled before RNA-Seq was performed on Illumina HiSeq4000 at the Vincent J. Coates Genomics Sequencing Laboratory at the California Institute for Quantitative Biosciences (QB3) at University of California, Berkeley. Libraries from LCM samples and STG blocks were sequenced to about 50 million 2x150bp reads per sample. For libraries prepared with the NuGEN Ovation SoLo kit a Custom R1 primer (NuGEN) was used in place of the standard Illumina forward read primer.

### Mapping, quantification of gene expression, and QC

RNA-seq reads were aligned to the GRCH37.p13 (hg19) reference genome via STAR (2.7.2a) using comprehensive gene annotations from GENCODE (v29 lifted over to hg19). Gene-level quantifications were calculated using featureCounts (v1.6.4), considering only uniquely-mapped reads. Quality control metrics were calculated using PicardTools (v2.21.2).

Gene-level counts were compiled and imported into R for downstream analyses. Expressed genes were defined as genes with non-zero count in at least 80% of samples. A total of 22,729 and 13,458 expressed genes from block tissue and LCM neurons, respectively, were used in the downstream analysis. Sample outliers were defined as samples with standardized sample network connectivity Z scores < -2 (81), and were removed.

A set of 105 RNA-Seq quality control metrics from the outputs of PicardTools (CollectAlignmentSummaryMetrics, CollectInsertSizeMetrics, CollectRnaSeqMetrics, CollectGcBiasMetrics, MarkDuplicates) were compiled for each group of samples (block tissue and LCM neurons). These measures were summarized by the top principal components (termed seqPCs), which explained a significant portion of the total variance of each dataset. These seqPCs were used as potential covariates for downstream analysis.

### Differential gene expression

Differential Gene Expression (DGE) analyses were performed using DESeq2 (1.22.2)(82) with default parameters. For block tissue data, diagnosis, sex, age, RNA integrity number (RIN), absorbance 260/280 ratio (A260/280) and top 3 seqPCs were used as covariates. For neuron data, diagnosis, sex, age, RNA library batch and top 3 seqPCs were used as covariates. To identify age-dependent differential expression, an interaction term between age and diagnosis was added to the above DESeq2 models.

### Differential alternative splicing

Local splicing analysis was performed using LeafCutter(83) as previously described(10). In brief, Clusters of variable spliced introns across all samples were called first. Then differential splicing between ASD and control group was identified in each data set (bulk tissue and neuron) by jointly modeling intron clusters using the Dirichlet-Multinomial generalized linear model (GLM). We controlled for the same covariates as above in the DGE analysis.

Intron clusters were first filtered to only keep clusters supported by at least 50 split reads across all samples, retaining introns of up to 100 kb and accounting for at least 1% of the total number of reads in the entire cluster. This intron count file was then used in the differential splicing (DS) analysis. For DS analysis, we further discarded introns that were not supported by at least one read in 5 or more samples. Clusters were then analyzed for DS if at least 3 samples in each comparison group (i.e. ASD or controls) had an overall coverage of 20 or more reads. P-values were corrected for multiple testing using the Benjamini-Hochberg (BH) method and used to select clusters with significant splicing differences (FDR < 0.1).

### Co-expression network analysis

Weighted gene co-expression network analysis (WGCNA)(23) defined modules of co-expressed genes from RNA-seq data. All covariates except for ASD diagnosis, sex and age were first regressed out from the expression datasets. The co-expression networks and modules were estimated using the blockwiseModules function with the following parameters: corType=bicorr; networkType=signed; pamRespectsDendro=F; mergeCutHeight=0.1, power=8, deepSplit=2, minModuleSize=40. Module eigengene/genotype associations were calculated using a linear model. Significance p-values were FDR-corrected to account for multiple comparisons. Genes within each module were prioritized based on their module membership (kME), defined as correlation to the module eigengene. For selected modules, the top hub genes were shown.

### Functional enrichment analysis

For co-expressed gene modules, enrichment for Gene Ontology (GO; Biological Process and Molecular Function) was performed using gProfileR R package(84). Background was restricted to the expressed set of genes. An ordered query was used, ranking genes by kME for WGCNA analyses.

For DGE, GO enrichment was performed using the GSEA algorithm as implemented in the clusterProfiler R package(85). All genes were ranked by log2 fold change.

Enrichment analyses were also performed using several established, hypothesis-driven gene sets including pre- and postsynaptic marker genes (77), genes with likely-gene-disruption (LGD) or LGD plus missense de novo mutations(DNMs) found in patients with neurodevelopmental disorders(78), genes with probability of loss-of-function intolerance (pLI) > 0.99 as reported by the Exome Aggregation Consortium (28), mutationally constrained genes(27), vulnerable ASD genes(79), CHD8 targets(29), FMRP targets (80), syndromic and highly ranked (1 and 2) genes from SFARI Gene database. Statistical enrichment analyses were performed using permutation test. One thousand simulated lists with similar number of genes, gene length distribution and GC-content distribution as the target gene list were generated, and the overlaps between each of the simulated list and the hypothesis-driven gene sets were calculated to form the null distribution. Significance p-value was calculated by comparing the actual overlap between target list and hypothesis-driven gene sets to the null distribution. All results were FDR-corrected for multiple comparisons.

### Ingenuity pathway analysis

We performed Ingenuity Pathway Analysis (IPA®, QIAGEN) to identify significantly over-represented pathways and to determine if they are activated or inhibited in ASD brain compared to control brain. IPA predicts the overall direction of the pathway (activation or inhibition) using a Z-score to statistically compare our datasets with expression patterns in the IPA knowledge base (86). This is achieved by considering the activation state of key molecules when the Pathway is activated and the molecules’ causal relationships. Z ≥ 2 signifies a pathway that is significantly activated, while Z ≤ −2 - significantly suppressed in ASD compared to control brain.

### Cell type enrichment analysis

Cell-type enrichment analysis for each co-expression module was performed using the Expression Weighted Cell Type Enrichment (EWCE) package in R(87). Cell type-specific gene expression data was obtained from single nucleus sequencing of adult human brains (88). The specificity metric of each gene for each cell type was computed as described(87). Enrichment was evaluated using bootstrapping. Z-score was estimated by the distance of the mean expression of the target gene set from the mean expression of bootstrapping replicates. P-values were corrected for multiple comparisons using FDR.

### GWAS enrichment analysis

The most recent ASD GWAS summary statistics were obtained from Grove *et al.*(30) Stratified LD score regression (sLDSC)(89) was used to test whether a gene set of interest is enriched for SNP-heritability in a given GWAS dataset. In brief, SNPs were assigned to custom gene categories if they fell within ±100 kb of any gene in a set. These categories were added to a full baseline model that includes 53 functional categories capturing a broad set of genomic annotations. The MHC region was excluded from all analyses. Enrichment was calculated as the proportion of SNP-heritability accounted for by each category divided by the proportion of total SNPs within the category. Significance was assessed using a block jackknife procedure, followed by Bonferroni correction for the number of gene sets.

## Data Availability

RNA-seq data will be submitted to the NCBI Sequence Read Archive (SRA), and accession code will be available before publication.

## Code Availability

All custom code used in this manuscript is available at https://github.com/gandallab/ASD_STG_LCM_RNAseq

## Acknowledgments

We are grateful to the families of our brain donors for their invaluable gift to autism research. We appreciate the contributions of Carolyn Komich Hare, clinical coordinator for BrainNet, for collection of the ADI-R as well as Robin Riedel, UC Davis, for figure design. Tissue was provided by the University of Maryland Brain and Tissue Bank, Dr. Daniel Campbell at Michigan State University, Autism Tissue Program (now Autism BrainNet, supported by the Simons Foundation), Brain Endowment for Autism Research Sciences (BEARS) at the UC Davis MIND Institute, and the Harvard Brain Tissue Resource Center. Laser capture microdissection was conducted using the CAMI core facility at UC Davis Center for Health and the Environment with instrumentation funding by NIHS10RR-023555. RNA sequencing was conducted using the Vincent J. Coates Genomics Sequencing Laboratory at UC Berkeley, supported by NIH S10 OD018174 Instrumentation Grant. This work is supported by NIH grants MH108909 (MPI Schumann, Stamova) and the Intellectual and Developmental Disabilities Research Center at UC Davis (1U54HD079125).

## Author Contributions

C.M.S., B.S. and B.P.A. conceived the project. C.M.S., B.S. and M.J.G. supervised the study. A.O., B.P.A., B.S. and C.M.S. designed and performed the experiments. P.Z., B.S. and M.J.G. designed and performed computational analyses. P.Z. wrote the paper with input from all co-authors.

