## Supplemental Figures for "Neuron-specific transcriptomic signatures indicate neuroinflammation and altered neuronal activity in ASD temporal cortex"

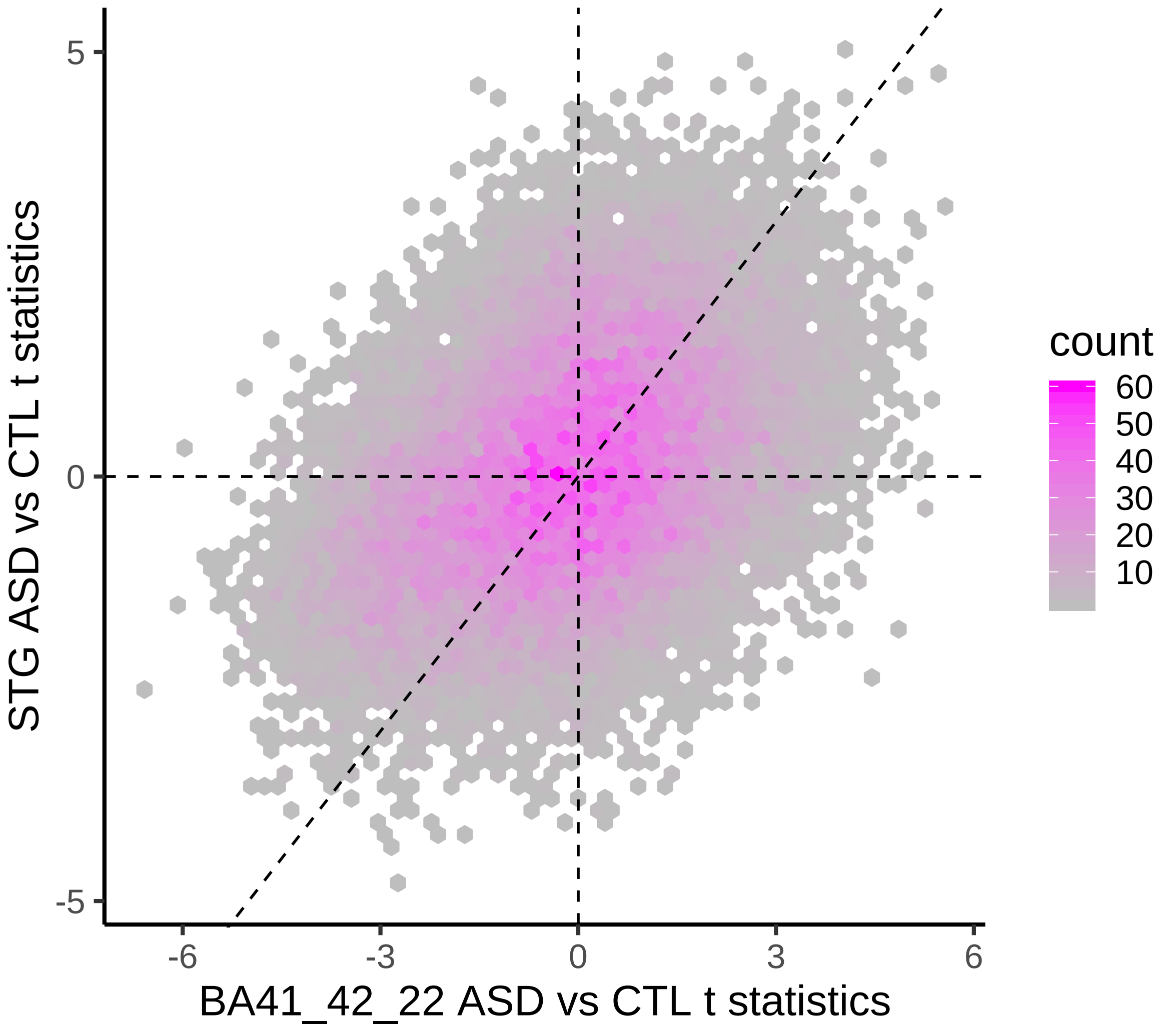


**Supplementary Figure 1**

Binned density scatter plot comparing the t-statistics for case versus control differential expression between this study and another study (ref. 12) comparing gene expression between ASD and controls in BA41, BA42, and BA22 bulk tissues; correlation between the statistics is 0.37 ($P < 10^{-16}$).

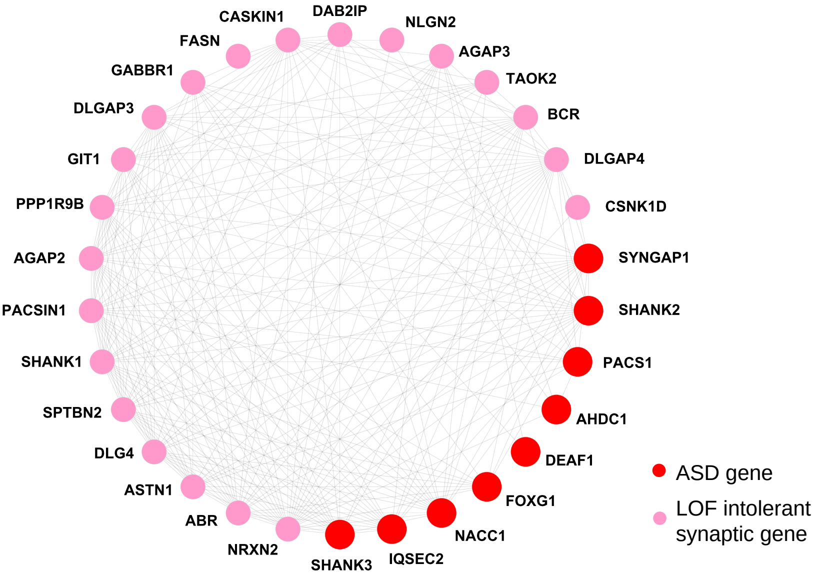


**Supplementary Figure 2**
Representative genes in module Block-M10. Known ASD risk genes were colored red. Synaptic genes that are intolerant to LOF mutation were colored pink.
Edges represent co-expression.

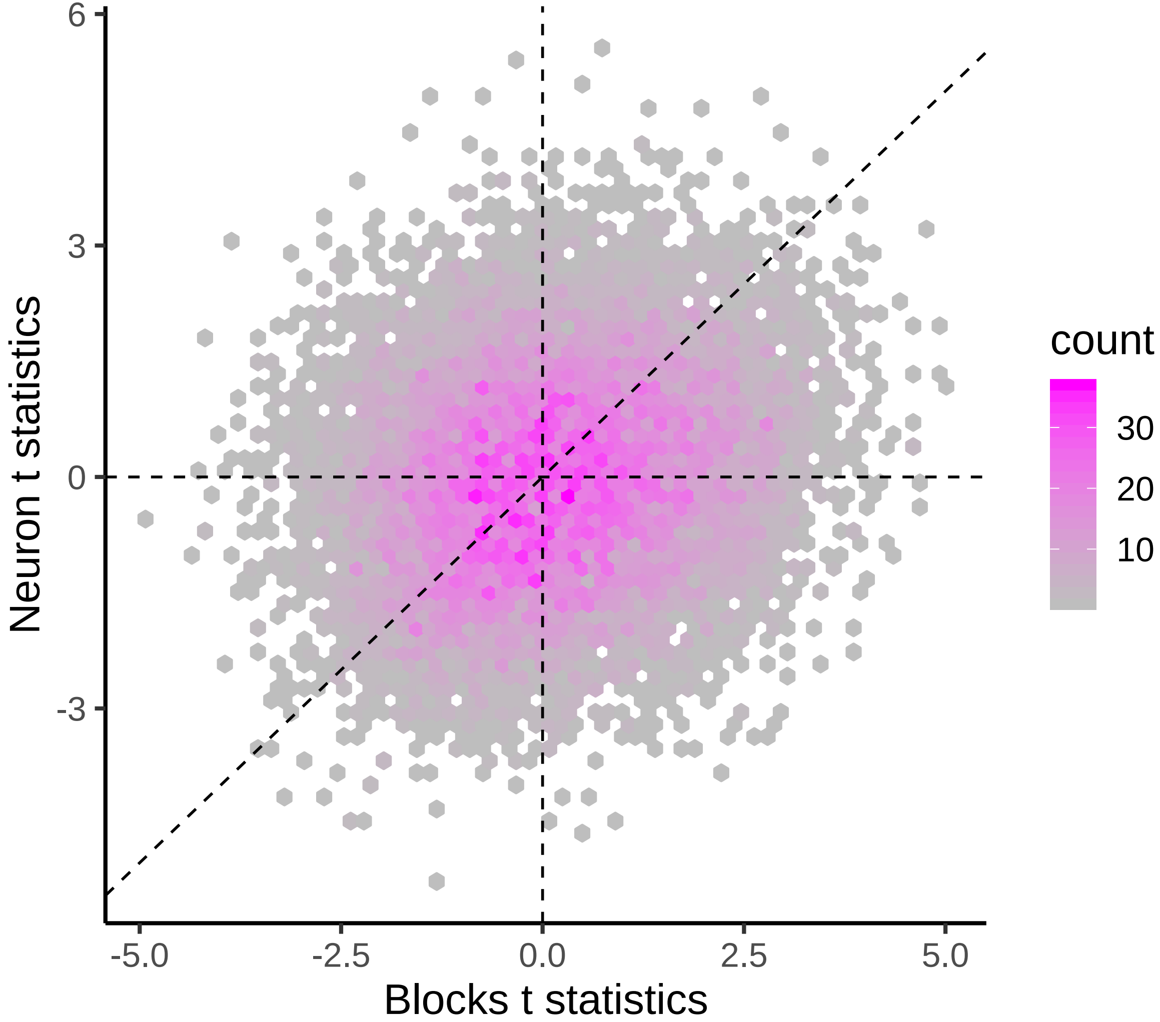


**Supplementary Figure 3**

Binned density scatter plot comparing the t-statistics for case versus control differential expression between neurons and bulk tissue; correlation between the statistics is 0.18 ($P < 10^{-16}$).

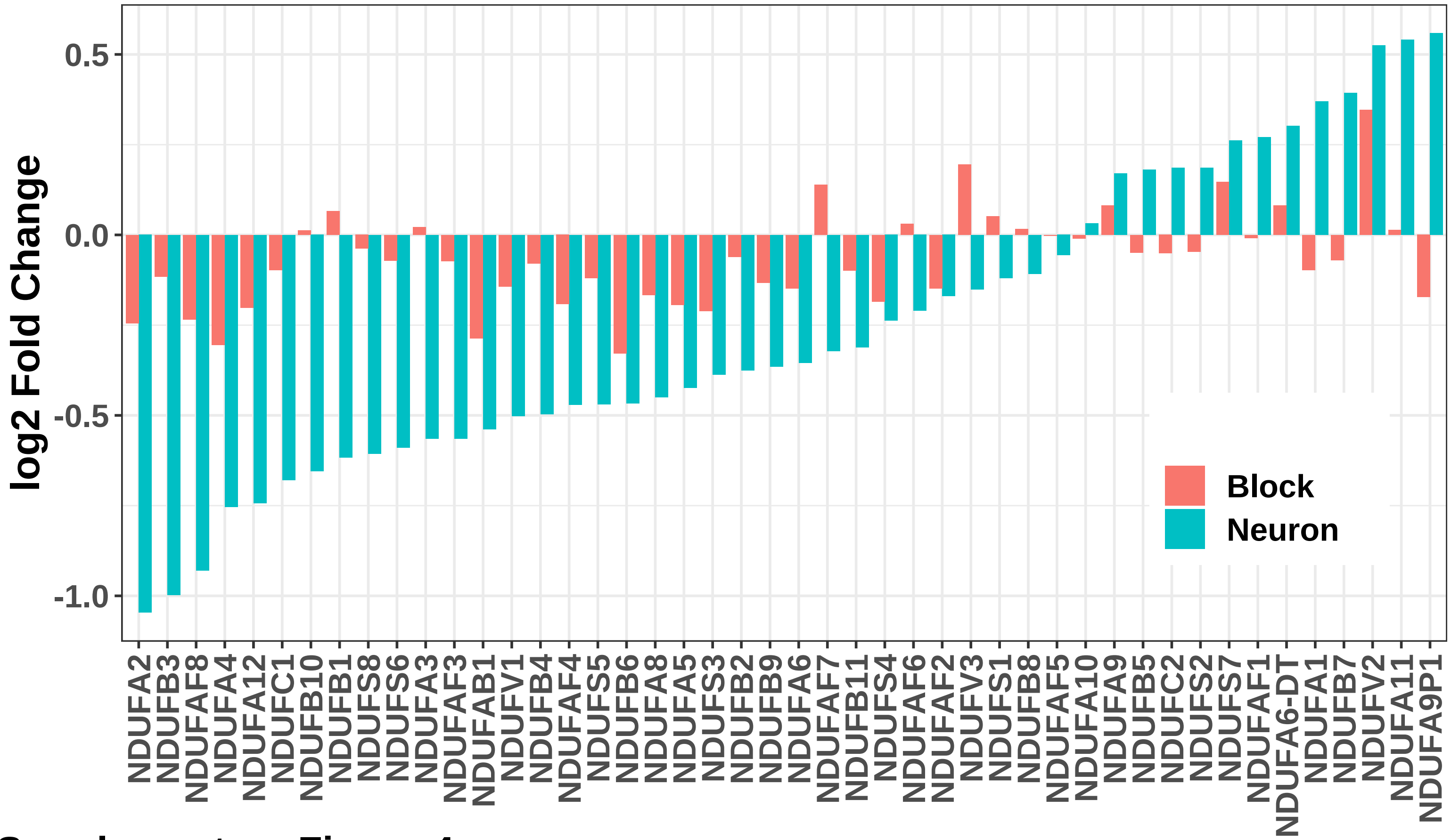


**Supplementary Figure 4**

Fold changes (ASD vs. CTL) of NADH:ubiquinone oxidoreductase (complex I) subunits in block tissue (red) and neurons (green).

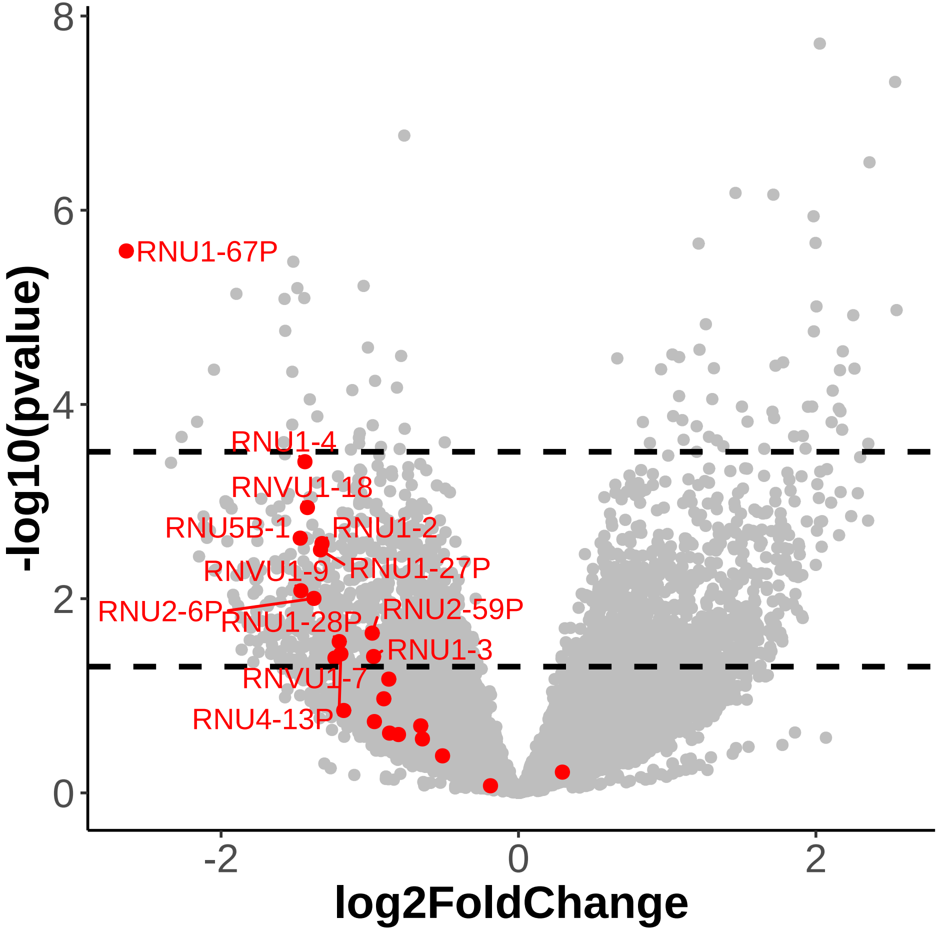


**Supplementary Figure 5**

Volcano plot showing differentially expressed genes in ASD neurons compared to control. snRNA genes were colored red.
